# Mini-Pcdh15b Gene Therapy Rescues Visual Deficits in a Zebrafish Model of Usher Syndrome Type 1F

**DOI:** 10.1101/2025.11.05.686814

**Authors:** Xinlan Chen, Daniel M Hathaway, Alex J Klein, Jennifer B Phillips, Kevin TA Booth, Muriel Heitsch, Anton Lytvyn, Bruce H Derfler, Jeremy Wegner, Corey Goldstein, Andrew Murphy, Sean G Megason, Monte Westerfield, David P Corey, Maryna V Ivanchenko

**Affiliations:** Department of Neurobiology, Harvard Medical School, Boston, Massachusetts, United States; Institute of Neuroscience, University of Oregon, Eugene, Oregon, United States; Department of Medical and Molecular Genetics, Indiana University School of Medicine, Indianapolis, Indiana, United States; Department of Systems Biology, Harvard Medical School, Boston, Massachusetts, United States

## Abstract

Usher syndrome type 1F (USH1F) is a severe inherited disorder caused by mutations in *PCDH15*, resulting in congenital deafness, vestibular dysfunction, and progressive retinal degeneration leading to blindness. While cochlear implantation can restore hearing, no therapeutic interventions currently exist for vision loss. Gene augmentation therapy represents a promising approach; however, the *PCDH15* coding sequence (∼5.3 kb) exceeds the packaging capacity of adeno-associated virus (AAV) vectors. To overcome this limitation, we previously engineered shortened “mini-PCDH15” constructs that retain key structural and functional domains while fitting within a single AAV. Among these, the mini-PCDH15-V4 variant successfully restored hearing in *Pcdh15*-deficient mice.

Here, we investigated the ability of a cone-targeted *mini-pcdh15b-V4* transgene to rescue vision in a zebrafish model of USH1F. Untreated *pcdh15b*-deficient zebrafish exhibited severe structural and functional defects of the retina, including disorganized and shortened photoreceptor outer segments, disrupted calyceal processes, and markedly reduced electroretinogram (ERG) and optokinetic response (OKR) performance. Targeted expression of mini-*pcdh15b*-V4 recapitulated typical localization of Pcdh15b to calyceal processes and outer segment membranes, rescued photoreceptor architecture, and re-established both structural organization and functional output. Treated mutants exhibited improved visual tracking behavior and full recovery of ERG a-wave and b-wave amplitudes, indicating restoration of photoreceptor and synaptic function. Importantly, mini-*pcdh15b*-V4 expression produced no adverse effects in wild-type or heterozygous fish, supporting the safety of cone-specific expression.

Together, these findings demonstrate that mini-*pcdh15b*-V4 can restore both photoreceptor structure and visual function in *pcdh15b*-deficient zebrafish. This work establishes the *pcdh15b* mutant zebrafish as a powerful preclinical model for studying USH1F retinopathy and supports the translational potential of rationally engineered mini-PCDH15 constructs as a feasible gene therapy approach for preventing or reversing vision loss in individuals with USH1F.

## INTRODUCTION

Usher syndrome (USH) is an autosomal recessive disorder characterized by sensorineural hearing loss, retinitis pigmentosa (RP), and vestibular dysfunction (Geleoc & El-Amraoui, 2020; Mathur & Yang, 2015). Of the three clinically defined subtypes of USH (USH1, USH2, USH3), Usher syndrome type 1 (USH1) is the most severe, presenting with congenital severe-to-profound deafness, vestibular areflexia at birth, and RP developing in the first decades of life (Mathur & Yang, 2015). Mutations in the protocadherin-15 gene (*PCDH15*) cause Usher syndrome 1F (USH1F), a genetic subtype of USH1 resulting from the loss or dysfunction of the adhesion protein, PCDH15 (Ahmed et al., 2006; Choudhary et al., 2020; De-la-Torre, Choudhary, Araya-Secchi, Narui, & Sotomayor, 2018; Dionne et al., 2018; Powers, Gaudet, & Sotomayor, 2017; Sotomayor, Weihofen, Gaudet, & Corey, 2012). In the neurosensory hair cells of the inner ear, PCDH15 interacts with cadherin 23 (CDH23) at their N termini to form tip links between adjacent stereocilia, essential for the transduction of mechanical stimuli into electrical receptor potentials. (Ahmed et al., 2001; Assad, Shepherd, & Corey, 1991; Corey, Akyuz, & Holt, 2019; Goodyear, Forge, Legan, & Richardson, 2010; Indzhykulian et al., 2013; Kazmierczak et al., 2007). In the visual system, the function of PCDH15 remains unclear. Studies of human and nonhuman primate photoreceptors have revealed the presence of PCDH15 along the calyceal processes, axially oriented microvillus-like structures that emanate from the inner segment and cup the base of the outer segment in both rods and cones (Sahly et al., 2012; Sharkova, Aparicio, Mouzaaber, Zolessi, & Hocking, 2024; Verschueren et al., 2022). Calyceal processes are essential for the formation of normally shaped photoreceptor outer segments (OS) and maintenance of normal photoreceptor function, and PCDH15*-*containing links are indispensable for calyceal process development and maintenance, as shown by knocking down *pcdh15* and *cdh23* in *Xenopus tropicalis* (Schietroma et al., 2017).

Therefore, the absence of PCDH15 in USH1F patients results in both profound congenital deafness, vestibular dysfunction and early-onset RP. Current treatment options for USH1F are limited to early cochlear implantation, which can restore hearing and facilitate communication, although patients may still face difficulties with speech perception in noisy environments (Davies et al., 2021). No approved therapies exist for RP and associated vision loss, or for the loss of vestibular function.

Delivery of the PCDH15 coding sequence to affected tissues using adeno-associated virus (AAV) could offer a therapeutic strategy, but the ∼5.3 kb coding sequence exceeds the packaging limit of AAV vectors (∼4.7 kb). We previously developed a rationally engineered “mini-PCDH15” (∼3.8 kb) by deleting selected extracellular cadherin (EC) repeats while preserving essential functional domains (Ivanchenko et al., 2023b). Among the engineered mini-PCDH15 variants tested, we identified one lacking five EC repeats, enabling the coding sequence to fit within a single AAV. When delivered to the cochleas of USH1F mouse models, this construct restored tip link formation, preserved hair bundle integrity, and rescued hearing (Ivanchenko et al., 2023b).

Although mouse models of USH1F have allowed us to study structural and functional auditory and vestibular defects associated with the disorder, retinal defects remain less well understood, as rodent and primate photoreceptor cells have major structural differences with rodents lacking calyceal processes (El-Amraoui & Petit, 2014). The lack of retinal degeneration or morphological defects in the USH1F mouse model may be due to these species differences (El-Amraoui & Petit, 2014; Sahly et al., 2012), or alternatively, it may be that mice do not live long enough to manifest retinal degeneration.

In contrast, zebrafish retain the full complement of vertebrate sensory systems, including taste, touch, smell, balance, vision, and hearing, making it a desirable model organism for studying vertebrate biology (Angueyra & Kindt, 2018; Moorman, 2001; Noel, Allison, MacDonald, & Hocking, 2022). Notably, zebrafish photoreceptors do possess calyceal processes, a key structural feature absent in rodents (Hodel et al., 2014; Miles, Blair, Emili, & Tropepe, 2021; Phillips et al., 2023; Tarboush, Chapman, & Connaughton, 2012). In zebrafish, PCDH15 is represented by two paralog genes, *pcdh15a* and *pcdh15b* (Seiler et al., 2005). Reduction of *pcdh15a* activity leads to deafness and vestibular defects, while visual defects are not evident, suggesting a role for *pcdh15a* in stereocilia morphology and function (Maeda, Pacentine, Erickson, & Nicolson, 2017). Conversely, mutations in *pcdh15b* cause early-onset and progressive visual defects resembling USH1F retinopathy (Miles et al., 2021; Phillips et al., 2023).

In this study, we established and characterized a zebrafish *pcdh15b* mutant model of USH1F retinopathy, which recapitulates key structural and functional features of the human USH1F, including disorganized and shortened photoreceptor outer segments (OS), defective calyceal processes, reduced light-evoked electrical activity, and impaired visual behavioral responses. Leveraging this model, we generated a stable transgenic zebrafish line expressing our cone-specific mini-Pcdh15b-V4 construct, previously shown to restore hearing in USH1F mice (Ivanchenko et al., 2023b). Targeted expression of mini-Pcdh15b-V4 in *pcdh15b*-deficient photoreceptors reproduced the subcellular localization of full-length Pcdh15b, preserved photoreceptor architecture, restored calyceal process organization, and rescued both visual behavior and cone-mediated retinal function. These results show that the *pcdh15b* mutant zebrafish is a powerful model for preclinical studies of USH1F and highlight mini-PCDH15 gene therapy as a promising approach for treating vision loss in USH1F.

## RESULTS

### Establishing a zebrafish model of pcdh15b-associated retinopathy for therapeutic testing

To test the efficacy of mini-PCDH15 gene therapy for blindness, we used a previously characterized zebrafish model of USH1F carrying a 7-bp deletion in exon 8 of the *pcdh15b* gene, which introduces a frameshift and premature stop codon (Phillips et al., 2023). Exon 8 was specifically targeted to disrupt all *pcdh15b* transcript variants (**Figure 1a**). The resulting mutation truncates the open reading frame, eliminating most of the extracellular cadherin repeats, as well as the transmembrane and cytoplasmic domains. Although earlier studies reported retinal abnormalities in *pcdh15b* mutants, additional vestibular defects—including circling, twirling swimming patterns, and impaired righting reflexes—have also been reported (Miles et al., 2021; Phillips et al., 2023; Seiler et al., 2005).

**Figure 1.**
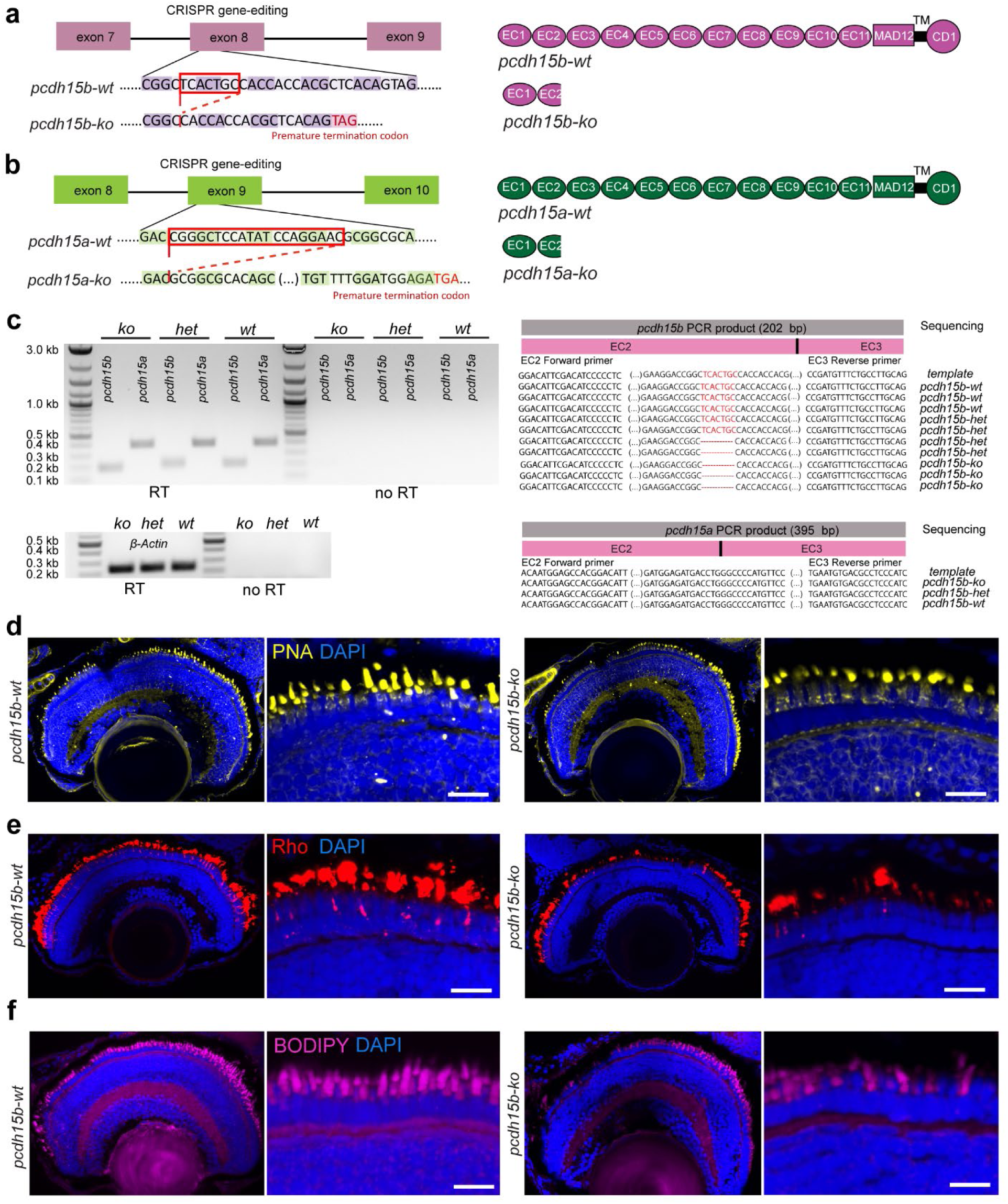
Zebrafish *Pcdh15b* mutants show abnormal retinal photoreceptor morphology. **a.** Schematic of *pcdh15b* zebrafish mutant generation with CRISPR/Cas9, resulting in a 7-bp deletion in exon 8 that introduced a frameshift and premature stop codon. **b.** Generation of the *pcdh15a* zebrafish mutant line. Schematic of *pcdh15a* zebrafish mutant generation with CRISPR/Cas9, resulting in a 20-bp deletion in exon 9 that introduced a frameshift and premature stop codon. The resulting mutation truncated the open reading frame, eliminating most of the extracellular cadherin repeats (EC), as well as a membrane-adjacent domain (MAD12), the transmembrane domain (TM), and a cytoplasmic domain (CD1). **c.** Agarose gel electrophoresis showing RT-PCR products of *pcdh15a* (395 bp) and *pcdh15b* (202 bp), with positive β-actin control (238bp). Expression of both *pcdh15a* and *pcdh15b* was detected in eyes across all genotypes of 7 dpf larvae (*pcdh15b-ko, pcdh15b-het, pcdh15b-wt*). Samples not treated with reverse transcriptase demonstrated no amplification. RT-PCR of beta-actin showed similar amplification levels among samples. Sequencing of retinal cDNA PCR products confirmed the presence of the *pcdh15b* mutation in *pcdh15b*-*ko* and *pcdh15b*-*het* samples, and its absence in wild-type controls (top). Sequencing of *pcdh15a* PCR products from the same retinal cDNA showed no detectable changes in the *pcdh15a* paralog (bottom). **d.** Representative confocal microscopy images of cone photoreceptor outer segments (OS) labeled with peanut agglutinin (PNA, yellow) demonstrated compromised OS morphology in 7-dpf *pcdh15b* knockout larvae (right) compared with wild-type controls (left). **e.** Representative images of rod photoreceptor OS labeled for rhodopsin (Rho, red) showed a reduced labeling of rods in 7-dpf *pcdh15b* knockout larvae (right) compared to wild-type controls (left). **f.** Representative confocal microscopy images of BODIPY labeling (magenta) of photoreceptor OS showed normal photoreceptor morphology in 7-dpf wild-type control larvae (left). In *pcdh15b* knockout larvae (right), the OS were shorter and more disorganized. Scale bars: **d-f** 10 µm

We examined *pcdh15a* and *pcdh15b* mRNA expression by reverse transcription PCR in homozygous mutant (*pcdh15b^−/−^;* hereafter referred to as *pcdh15b-ko),* heterozygous (*pcdh15b^+/−^; pcdh15b-het*), and wild-type (*pcdh15b^+/+^; pcdh15b-wt*) zebrafish at 7 days post-fertilization (dpf) and detected expression of both genes in the eye in all genotypes (**Figure 1c**). Sanger sequencing confirmed the presence of the *pcdh15b* mutation in knockout and heterozygous samples but not in wild-type samples, further validating the mutant line used in this study (**Figure 1c**).

To assess the retinopathy phenotype in larval zebrafish, we implemented a 72-hour light exposure protocol designed to exacerbate the retinopathy. Larvae maintained in transparent Petri dishes were transferred at 3 dpf to a custom incubator equipped with an overhead LED light source, where they were exposed to a controlled 12-hour light/12-hour dark cycle at an intensity of 3000 lux. This approach has been used in prior studies, which have demonstrated that environmental light exposure can modulate the severity of retinal phenotypes in USH models. Specifically, *pcdh15b* mutant zebrafish exhibit exacerbated photoreceptor structural defects and increased cell death under bright light conditions, whereas darkness is protective (Miles et al., 2021; Phillips et al., 2023).

Histological analysis of retinal cryosections from *pcdh15b* mutant and wild-type larvae at 7 dpf revealed significant disruptions in photoreceptor morphology in mutants, consistent with previous reports (Miles et al., 2021; Phillips et al., 2023). To further characterize these defects, we labeled the retinas with various markers for rod and cone photoreceptor OS. Staining with peanut agglutinin lectin (PNA), a marker for cone photoreceptor OS, showed a clear reduction in both the size and number of cone OS in *pcdh15b* mutants at 7 dpf (**Figure 1d**). Similarly, staining for rhodopsin, a marker for rod photoreceptor OS, was reduced, particularly in the central region of the retina in *pcdh15b* mutants at 7 dpf (**Figure 1e**). Additionally, we used BODIPY to label lipid membranes, which are particularly dense in photoreceptor OS. In *pcdh15b* knockout larvae, BODIPY staining was noticeably reduced in the peripheral, ventral, and central regions of the retina. The OS of cone photoreceptors appeared disorganized in the mutants, and the remaining OS were, on average, shorter compared to those in control siblings, likely due to the abnormal morphology of the OS (**Figure 1f**).

To determine Pcdh15 protein expression in the retina of 7 dpf zebrafish larvae, we tested commercially available antibodies targeting the extracellular region of human PCDH15 (EC1-EC11) that recognize both *pcdh15b* and *pcdh15a* isoforms. To assess photoreceptor morphology, we co-labeled the retinas with BODIPY to mark photoreceptors, and with phalloidin to highlight actin-rich calyceal processes, which normally form a barrel-like fringe around the OS and are thought to contribute to disc morphogenesis and structural stabilization of OS (**Figure 2a**).

**Figure 2.**
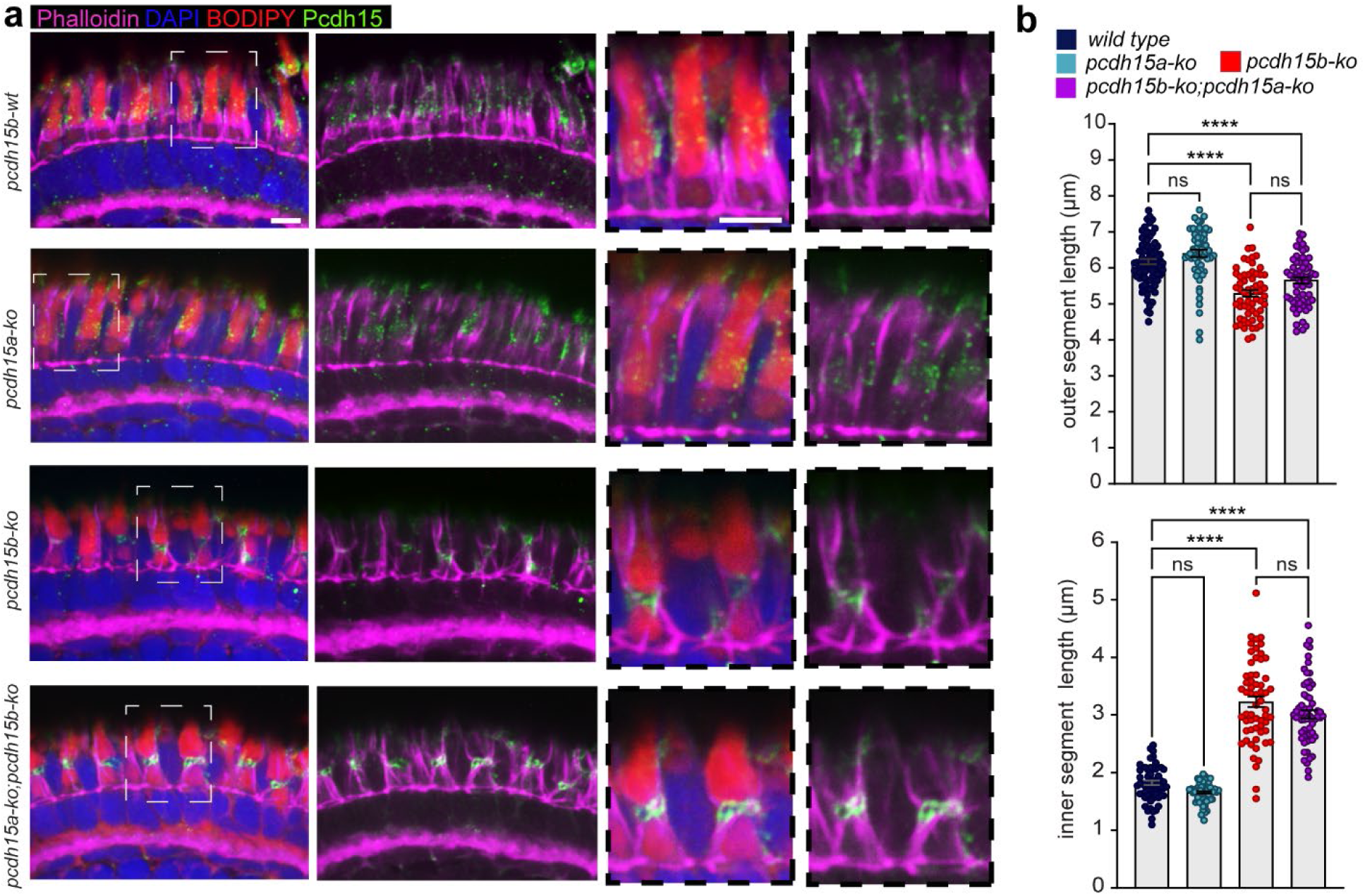
Zebrafish *Pcdh15b* mutants show reduced Pcdh15 protein expression in the retina. **a.** Representative images of photoreceptor calyceal processes *(phalloidin, magenta) and* Pcdh15 protein localization *(green*). In *pcdh15b-wt* and *pcdh15a-ko* larvae (top panels), Pcdh15 is prominently localized along calyceal processes, within photoreceptor OS, and as puncta in the presumptive synaptic region. In contrast, Pcdh15 expression is absent from calyceal processes in *pcdh15b-ko* and *pcdh15a-ko*;*pcdh15b-ko* double mutants (bottom panels). Additionally, *pcdh15b-ko* mutants and *pcdh15a-ko*;*pcdh15b-ko* double mutants show abnormal accumulation of Pcdh15 in the ellipsoid region, forming a gap between inner segments (IS) and OS. Photoreceptor OS membranes are labeled with BODIPY (red), and nuclei with DAPI (blue). **b.** Quantification of OS (top) and IS (bottom) lengths in *pcdh15b-wt*, *pcdh15a-ko, pcdh15b-ko,* and double-mutant (*pcdh15a-ko;pcdh15b-ko*) zebrafish larvae at 7 dpf. Data are shown as individual measurements with mean ± SEM. ****p < 0.0001; ns, not significant; one-way ANOVA with Šidák’s multiple comparisons test. Scale bars: 5 µm

In wild-type larvae, Pcdh15 was detected prominently along calyceal processes, within the photoreceptor OS, and as punctate staining within the presumptive synaptic region. In *pcdh15b-ko* larvae, Pcdh15 labeling was markedly reduced along calyceal processes and in the synaptic region. However, residual signal remained at the inner segment–outer segment junction within the ellipsoid region, likely reflecting either expression of *pcdh15a* isoforms or accumulation in the endoplasmic reticulum of truncated Pcdh15b protein corresponding to the first two extracellular domains.

To investigate this further, we generated *pcdh15a* and *pcdh15b* double mutants by crossing heterozygous animals for each gene. Previous studies of the *pcdh15a* ‘orbiter’ mutant suggest that *pcdh15a* is essential for stereocilia morphology and inner ear hair cell function but does not contribute to retinal phenotypes (Seiler et al., 2005). Our *pcdh15a* mutant carries a 20-bp deletion in exon 9, resulting in a frameshift and premature stop codon that disrupts all *pcdh15a* transcript variants (**Figure 1b**). This truncation eliminates most of the extracellular cadherin repeats along with the transmembrane and cytoplasmic domains.

Next, we further analyzed *pcdh15a-ko* and double mutant combinations by immunostaining (**Figure 2a**). In *pcdh15a-ko* larvae, Pcdh15 expression and photoreceptor morphology appeared largely similar to that in wild-type larvae. In contrast, both *pcdh15b-ko* and *pcdh15a-ko*;*pcdh15b-ko* (double mutant) larvae showed strongly reduced Pcdh15 labeling along calyceal processes. In these mutants, a residual signal was again detected within the ellipsoid zone, corresponding to the region containing the endoplasmic reticulum. The detected signal likely corresponds to translated fragments of *pcdh15b* containing the first two extracellular domains, which contain the antibody epitope.

Phalloidin staining of calyceal processes revealed pronounced defects in retinas from both *pcdh15b-ko* and double mutant larvae. The calyceal processes appeared fused, splayed, and branched, indicating disrupted cytoskeletal organization (**Figure 2a**). Additionally, these double mutants exhibited shortened photoreceptor OS. In contrast, *pcdh15a-ko* larvae exhibited calyceal process and photoreceptor organization comparable to that of wild-type siblings.

Quantification of photoreceptor morphology revealed significant shortening of the OS in *pcdh15b-ko* and double-mutant (*pcdh15a-ko;pcdh15b-ko*) larvae compared to wild-type and *pcdh15a-ko* siblings (**Figure 2b**). In contrast, inner segments (IS) were significantly longer in *pcdh15b-ko* and double mutants but remained unaffected in *pcdh15a-ko* larvae. This elongation likely reflects an expansion of the ellipsoid region and partial detachment of the OS.

Together, these findings highlight the essential role of *pcdh15b* in maintaining normal photoreceptor structure and support the idea that the *pcdh15b* paralog is the primary contributor to Pcdh15 function in the zebrafish retina, with *pcdh15a* providing only limited compensatory activity.

### Generation and characterization of a stable transgenic zebrafish line expressing mini-V4

To generate functional Pcdh15 constructs compatible with AAV packaging limits, we leveraged prior atomistic structural analyses and functional assessments of individual EC repeats. PCDH15 contains 11 EC repeats with conserved Ca²⁺-binding residues critical for force transmission, but not all repeats are essential for function. By removing 3–5 non-essential repeats, we produced mini-PCDH15 variants 25–40% shorter than the full-length protein. Among these designs, mini-HA.PCDH15-V4 retains EC1–3 and EC9–11, followed by MAD12, the transmembrane domain and the intracellular CD1 domain, preserving key functional domains while reducing overall size (**Figure 3a**). We first tested mini-PCDH15 variants in the inner ears of mouse models of USH1F deafness, where mini-V4 successfully restored hearing (Ivanchenko et al., 2023a). Based on this demonstrated efficacy in mouse cochlea, we next generated a stable transgenic zebrafish line expressing zebrafish mini-HA.Pcdh15b-V4 (hereinafter referred to as mini-V4) to test its ability to rescue retinal phenotypes associated with *pcdh15b* loss.

**Figure 3:**
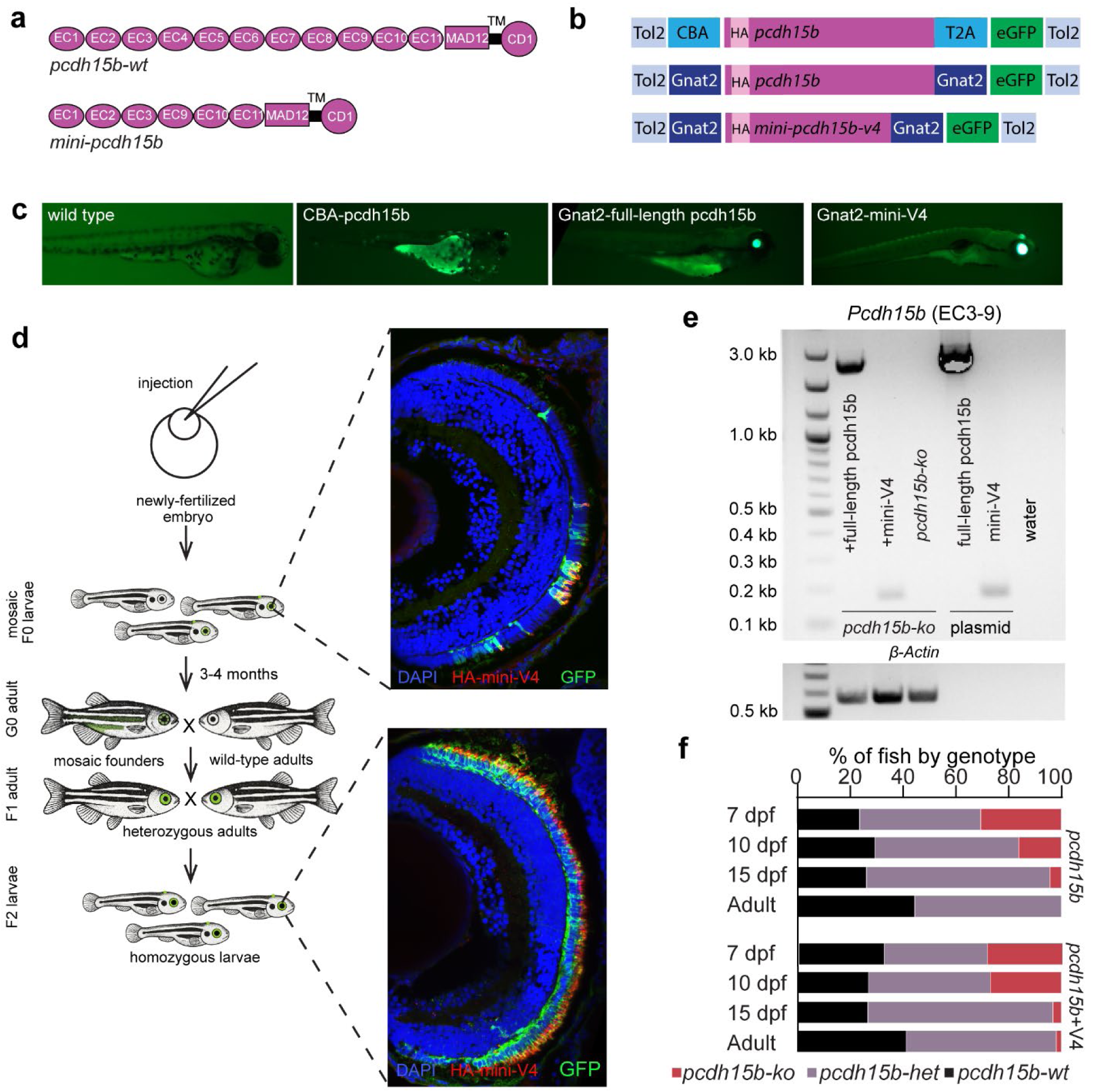
A stable *pcdh15b* mutant line expressing mini-V4 demonstrates proper localization along calyceal processes, within photoreceptor OS, and as puncta in the synaptic region. **a.** Schematic representation of Pcdh15b. The top diagram shows the domain structure of the full-length wild-type Pcdh15b protein, which includes 11 extracellular cadherin (EC) repeats, a membrane-adjacent domain (MAD12), the transmembrane domain (TM), and a cytoplasmic domain (CD1). The bottom diagram depicts mini-Pcdh15b-v4, a shorter version designed to preserve key functional domains, which retains EC1-3 and EC9-11, followed by MAD12, the TM, and CD1. **b.** Tol2 transposon-bicistronic donor plasmid designs to introduce full-length*-pcdh15b* tagged with human influenza hemagglutinin (HA) or *mini-V4* and *eGFP* driven by CBA or Gnat2 promoters. **c** Representative fluorescence images of 4-6 dpf zebrafish larvae expressing *pcdh15b* transgenes. No fluorescence was observed in wild-type (*pcdh15b-wt*) larvae. Broad and ectopic expression was seen in larvae expressing *pcdh15b* under the CBA promoter, which causes developmental malformations in F0 embryos. In contrast, expression under the cone-specific Gnat2 promoter of full-length *pcdh15b* or *mini-V4* yielded expression largely confined to the retina and pineal gland, without developmental abnormalities. **d.** Generation of the stable transgenic zebrafish line. At the one-cell stage, fertilized zebrafish eggs were injected with a mixture of the Tol2 transposon-donor plasmid and Tol2 transposase mRNA. Due to the random nature of integration, the injected F0 generation has mosaic expression of mini-V4 and eGFP. Transgenic fish with integration in germ cells passed the transgene to the F1 generation, and F2 larvae displayed stable transgene expression. **e** Genomic DNA analysis of *pcdh15b* expression using primers targeting EC3–EC9. A strong band was observed in *pcdh15b-ko* larvae injected with *mini-V4* and in *pcdh15b-ko* larvae co-injected with full-length *pcdh15b*, confirming transgene presence. The band size matched that of the plasmid-only controls. No amplification was detected in *pcdh15b-ko* larvae lacking transgene injection. β-Actin served as a loading control. **f** Genotypic distribution (*pcdh15b-ko, -het, and -wt*) at different developmental stages (7, 10, 15 dpf, and adult) in zebrafish with native *pcdh15b* or *pcdh15b* mutants expressing *mini-V4*. Longitudinal analysis revealed a significant reduction in the ratio of *pcdh15b-ko* larvae in breeding clutches starting around 10 dpf, and they were almost completely gone by 15 dpf. The *pcdh15b-ko* larvae with *mini-V4* began to die around 15 dpf and rarely reached the adult stage.

First, Tol2 transposon-based bicistronic donor plasmids were constructed to drive expression, under control of the ubiquitous CBA promoter, of full-length Pcdh15b and a GFP reporter separated by a T2A self-cleaving peptide (**Figure 3b**). However, microinjection of these constructs into one-cell stage embryos together with Tol2 transposase mRNA resulted in developmental malformations in F0 embryos, possibly due to aberrant cell-cell adhesion caused by widespread ectopic expression of Pcdh15b (**Figure 3c**). To overcome this, we next employed a cone photoreceptor-specific promoter (Gnat2) to restrict expression to the retina (Kennedy et al., 2007; Morrissey, Shelton, Brockerhoff, Hurley, & Kennedy, 2011; Smyth, Di Lorenzo, & Kennedy, 2008). We constructed Tol2-based bicistronic donor plasmids carrying either full-length *HA.Pcdh15b* or *mini-V4*, along with a GFP reporter, each driven by a Gnat2 promoter (**Figure 3b**). Microinjection of these constructs into one-cell stage embryos, together with Tol2 transposase mRNA, resulted in normal embryonic development and mosaic expression in F0 fish. Stable germline transmission was confirmed in F1 adults, and homozygous F2 larvae displayed robust transgene expression in the retina as visualized by GFP (**Figure 3c-e**).

To assess survival across genotypes, we genotyped multiple clutches from heterozygous crosses of both native *pcdh15b* mutants and *pcdh15b* mutants carrying a genomically integrated *mini-V4* transgene at 7, 10, and 15 dpf, as well as in adulthood. Homozygous *pcdh15b-ko* larvae progressively declined as a proportion of the total starting around 10 dpf and were nearly absent by 15 dpf, indicating a significant survival deficit (**Figure 3f**), suggesting that loss of Pcdh15b function in the retina and in other tissues such as the brain or vestibular organs, where the gene is also expressed, contributes to lethality once yolk reserves are exhausted and active feeding begins (Phillips et al., 2023; Seiler et al., 2005).

A similar pattern was observed in *pcdh15b-ko+mini-V4* animals; however, survival appeared slightly improved compared to native knockouts (**Figure 3f)**. One of these animals reached adult stage. The survivor had swimming remained impaired; it did not circle continuously but showed intermittent imbalance. Because mini-V4 expression is driven by a photoreceptor-specific promoter, we consider extra-retinal rescue of vestibular function unlikely; rather, partial visual restoration likely improves feeding efficiency sufficiently to permit survival in a minority of animals. All genotypes were present at expected Mendelian ratios at 7 dpf, indicating normal early development.

### Mini-V4 restores photoreceptor morphology in *pcdh15b* mutant zebrafish

To determine whether expression of mini-V4 can rescue retinal defects in *pcdh15b* mutants, we analyzed photoreceptor morphology at 7 dpf in *pcdh15b-wt, pcdh15b-het* and *pcdh15b-ko* larvae, either carrying the *mini-V4* transgene or not (**Figure 4**). Again, BODIPY (magenta) labeled the membranes of inner and outer segments, and phalloidin (green) labeled the actin of calyceal processes (**Figure 4a**). In *pcdh15b-ko* larvae, photoreceptors showed disorganized and shortened OS, and an enlarged gap between the IS and OS. Actin-rich calyceal processes appeared abnormally fused, splayed, and mislocalized around the OS. We noted, first, that the pronounced reduction in total photoreceptor number observed in *pcdh15b-ko* larvae was completely reversed by mini-V4 expression (**Figure 4a, b**), indicating that mini-V4 can function to enhance cell survival or maintenance.

**Figure 4:**
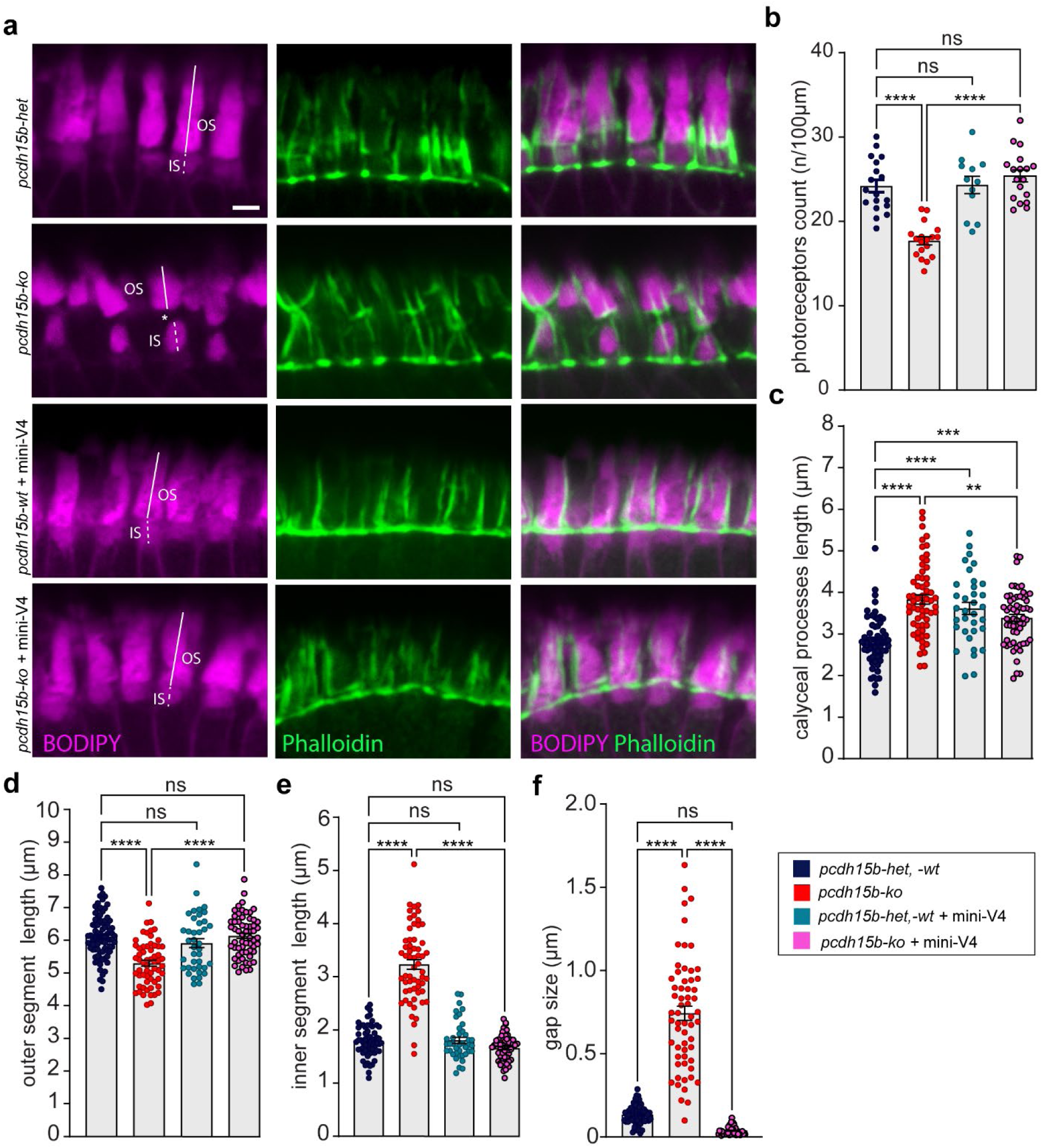
Photoreceptor disorganization in *pcdh15b* mutants is rescued in mutants that expressed mini-V4. **a** Confocal images of photoreceptors in 7 dpf *pcdh15b-het*, *pcdh15b-ko*, and *pcdh15b-het* or *pcdh15b-ko* animals expressing mini-V4. In *pcdh15b-ko* larvae, photoreceptors showed disorganized and shortened OS, and an enlarged gap between the BODIPY-labeled IS and OS (magenta). Calyceal processes labeled with phalloidin (green) appeared abnormally fused, splayed, and mislocalized around the OS. Expression of mini-V4 restored the length and alignment of photoreceptor segments and rescued the structural integrity of calyceal processes. Note the abnormal OS detachment and disc disintegration (holes) in *pcdh15b-ko*, which were rescued by mini-V4. **b** Quantification of photoreceptor number per 100 μm of retina length across genotypes. **c-f** Quantification of calyceal process length (**c**), OS length (**d**), IS length (**e**), and IS-OS gap size (**f**) across genotypes. Expression of mini-V4 in both heterozygous and homozygous backgrounds significantly improved all parameters compared to untreated *pcdh15b*-*ko*. Data are shown as individual measurements with mean ± SEM. * < 0.05; ** p < 0.01; *** p < 0.001; **** p < 0.0001; ns, not significant; one-way ANOVA with Šidák’s multiple comparisons test. Scale bar: 2.5 μm.

Next, we quantified the length of calyceal processes in *pcdh15b-ko* larvae (**Figure 4a, c**). The analysis revealed a significant increase in calyceal process length in the *pcdh15b-ko* group compared to *pcdh15b-wt* controls, suggesting dysregulated actin dynamics in the absence of *pcdh15b*. Expression of mini-V4 in both heterozygous and homozygous backgrounds resulted in an increase in calyceal process length, exceeding the wild-type levels.

In *pcdh15b-ko* animals, confocal imaging revealed severely disorganized photoreceptors with shortened OS (**Figure 4a, d**), longer IS (**Figure 4a, e**), an enlarged gap between IS and OS (**Figure 4a, f**), and signs of OS instability, including partial detachment and disc disintegration. In contrast, expression of mini-V4 in homozygous mutant animals markedly improved photoreceptor architecture. Treated *pcdh15b-ko* larvae exhibited restoration of OS and IS length, normalization of IS–OS spacing, and proper organization of calyceal processes (**Figure 4a, c–f**). Notably, photoreceptors of heterozygous animals expressing mini-V4 did not show structural abnormalities, apart from a modest increase in calyceal process length.

### Mini-V4 expression restores photoreceptor ultrastructure in *pcdh15b* mutant zebrafish

To assess the impact of *pcdh15b* loss on photoreceptor ultrastructure, we performed transmission electron microscopy and scanning electron microscopy on retinas from 7 dpf zebrafish larvae (**Figure 5** and **Figure 6**).

**Figure 5:**
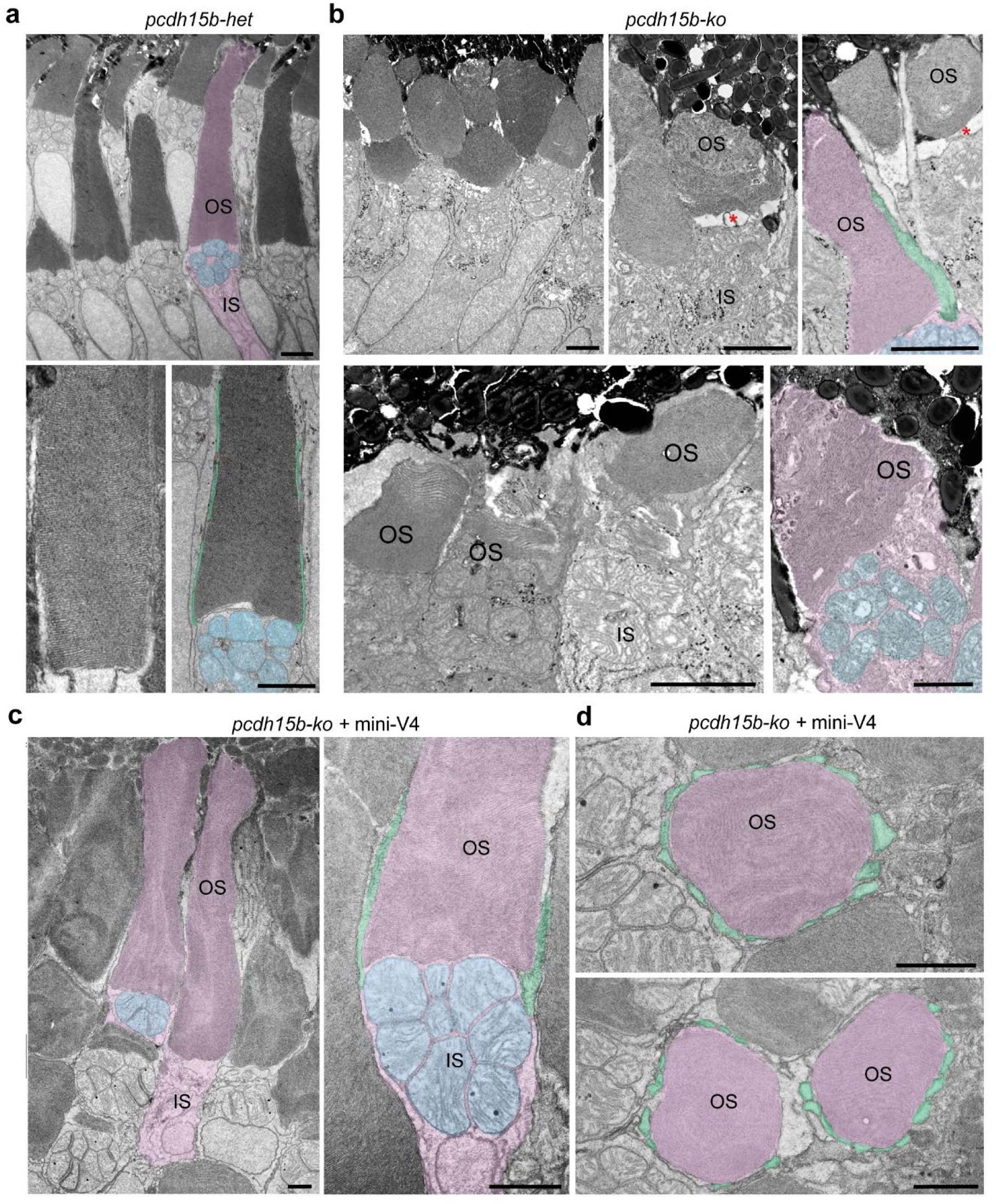
pcdh15b mutant photoreceptors show abnormal morphology. Color overlays: photoreceptor (purple), mitochondria (blue), and calyceal processes (green). **a** Representative transmission electron micrographs in 7 dpf *pcdh15b-het* larvae showed normal photoreceptor morphology with organized and elongated OS, connected IS, and thin calyceal processes aligned along the OS. **b** Representative transmission electron micrographs in *7 dpf pcdh15b-ko* larvae displayed disorganized photoreceptor morphology. OS were often disorganized or detached (red asterisks), with fragmentation, swelling, and irregular disc membranes. Calyceal processes (green) were reduced or absent, and the alignment between OS and IS was disrupted. **c** In *pcdh15b-ko* larvae expressing mini-V4, photoreceptor structure was substantially improved. OS and IS compartments appeared normal in morphology and alignment, with reestablished calyceal processes (green) and preservation of OS discs, resembling the heterozygous condition. **d** Representative TEM images of horizontal cross-sections through the OS in *pcdh15b-ko* larvae expressing mini-V4. Scale bars: 1 µm.

**Figure 6:**
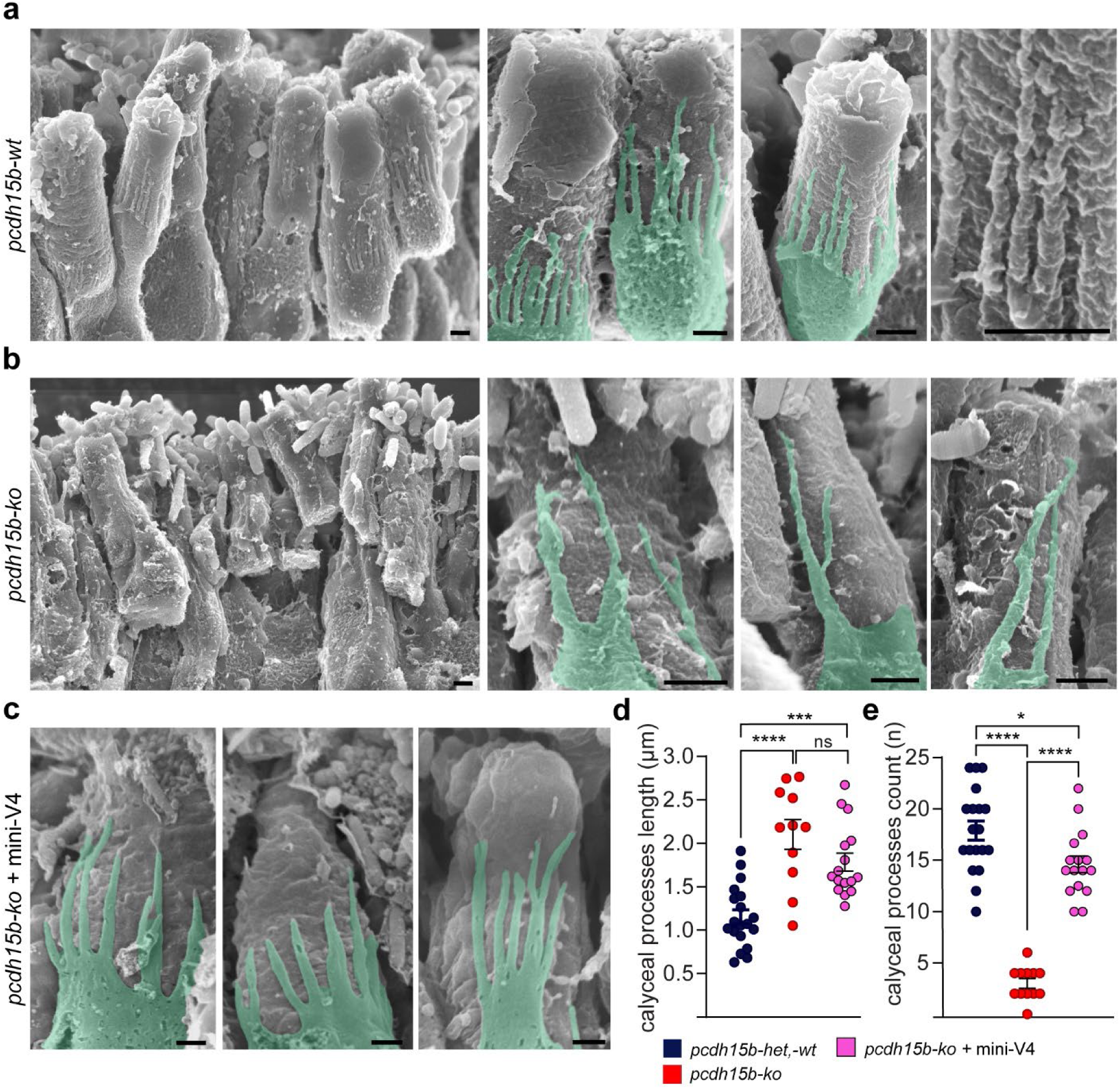
Mini-V4 expression restores both morphology and quantity of calyceal processes in *pcdh15b* mutant larvae. a–c. Scanning electron microscopy images of photoreceptor OS in 7 dpf zebrafish larvae. **a** Control *pcdh15b-wt* retinas displayed well-organized photoreceptors with calyceal processes (color overlay in green) surrounding the OS membrane. **b** *pcdh15b-ko* larvae showed a marked reduction in intact calyceal processes surrounding the OS compared to wild-type siblings, with the remaining processes appearing abnormally thick and branched. **c** Expression of mini-V4 in *pcdh15b-ko* larvae restored calyceal process morphology, resulting in elongated and well-organized structures surrounding the OS, resembling those observed in wild-type siblings. **d** Quantification of calyceal process length quantification, measured from the branching point of the protrusions, shows significantly longer calyceal processes in *pcdh15b-ko* and *pcdh15b-ko* with mini-V4 expression compared to *pcdh15b-wt*. **e** Quantification of calyceal processes counts per photoreceptor demonstrates a reduced number of calyceal processes in *pcdh15b-ko* that was restored with mini-V4 expression. Data are shown as individual measurements with mean ± SEM. * < 0.05; ** p < 0.01; *** p < 0.001; **** p < 0.0001; ns, not significant; one-way ANOVA with Šidák’s multiple comparisons test. Scale bar: **a-c** 0.5 μm.

Transmission electron microscopy of *pcdh15b* heterozygous control photoreceptors revealed normal morphology, characterized by elongated, tightly stacked OS discs; well-aligned IS enriched with mitochondria; and slender calyceal processes extending along the base of the OS (**Figure 5a**). In contrast, *pcdh15b*-deficient photoreceptors showed severe structural abnormalities. OS were frequently disorganized, shortened, or detached from the IS, with fragmented or swollen disc membranes. Calyceal processes were notably reduced or absent, and the alignment between OS and IS was disrupted (**Figure 5b**). Consistent with these observations, previous ultrastructural studies (Miles et al., 2021) reported that *pcdh15b* mutant retinas exhibit abnormal directional growth of OS discs, leading to misshapen and shortened OSs. These defects were evident in larval photoreceptors, with cones appearing more severely affected than rods (Miles et al., 2021), suggesting also a developmental rather than degenerative origin. Notably, we found that the expression of mini-V4 in *pcdh15b-ko* larvae markedly improved photoreceptor architecture (**Figure 5c** and **d**). The overall morphology of OS and IS appeared restored, with reformation of calyceal processes and preservation of disc organization, closely resembling the structure observed in heterozygous control siblings.

To examine photoreceptor ultrastructure further, we performed scanning electron microscopy on retinal tissue and noticed a reduction in intact calyceal processes surrounding the OS in *pcdh15b-ko* mutants compared to wild-type siblings. In *pcdh15b-wt* retinas, photoreceptors displayed well-organized OS surrounded by thin calyceal processes arranged in a uniform ring at the OS base (**Figure 6a**). In contrast, *pcdh15b-ko* larvae exhibited many fewer calyceal processes per photoreceptor, and the remaining structures appeared abnormally thick and branched (**Figure 6b**). Expression of mini-V4 in *pcdh15b-ko* larvae fully restored calyceal process morphology, producing elongated, well-aligned structures that closely resembled those of wild-type siblings (**Figure 6c**).

Quantitative analysis confirmed these observations. Calyceal process length, measured from the branching point of each protrusion, was significantly greater in *pcdh15b-ko* larvae compared to wild-type controls, whereas the number of calyceal processes per photoreceptor was significantly reduced in *pcdh15b-ko* larvae (**Figure 6d**). In mini-V4–expressing mutants, calyceal processes remained elongated, but their number was restored to levels comparable to wild-type (**Figure 6e**). These findings indicate that mini-V4 not only corrects the morphological abnormalities of calyceal processes in *pcdh15b* mutants but also restores their number, supporting a role in maintaining the structural integrity of these photoreceptor support structures.

### Targeted expression of mini-V4 recapitulates full-length Pcdh15b localization in *pcdh15b*-knockout retina

To determine whether mini-V4 retains the subcellular localization of full-length Pcdh15b, we examined transgenic *pcdh15b-ko* and *pcdh15b-het* retinas expressing HA-tagged mini-V4 (HA.mini-V4) or full-length Pcdh15b. Confocal imaging at 7 dpf revealed that both mini and full-length constructs were enriched at the apical region of photoreceptors, overlapping BODIPY-labeled OS (**Figure 7a**). In *pcdh15b*-*ko* larvae, HA.mini-V4 and full-length Pcdh15b displayed indistinguishable localization patterns, with robust anti-HA signal along the calyceal processes and OS membranes. Similar localization was observed in *pcdh15b-het* larvae expressing HA.mini-V4 **(Figure 7a,b** and **Supplementary Figure 1).** Higher-magnification imaging confirmed punctate HA signal along the microvilli-like calyceal processes and at the bases of the OS (**Figure 7b**), closely matching the pattern of full-length Pcdh15b. This distribution was consistent in both cross-sectional and in face views, confirming proper membrane targeting.

**Figure 7:**
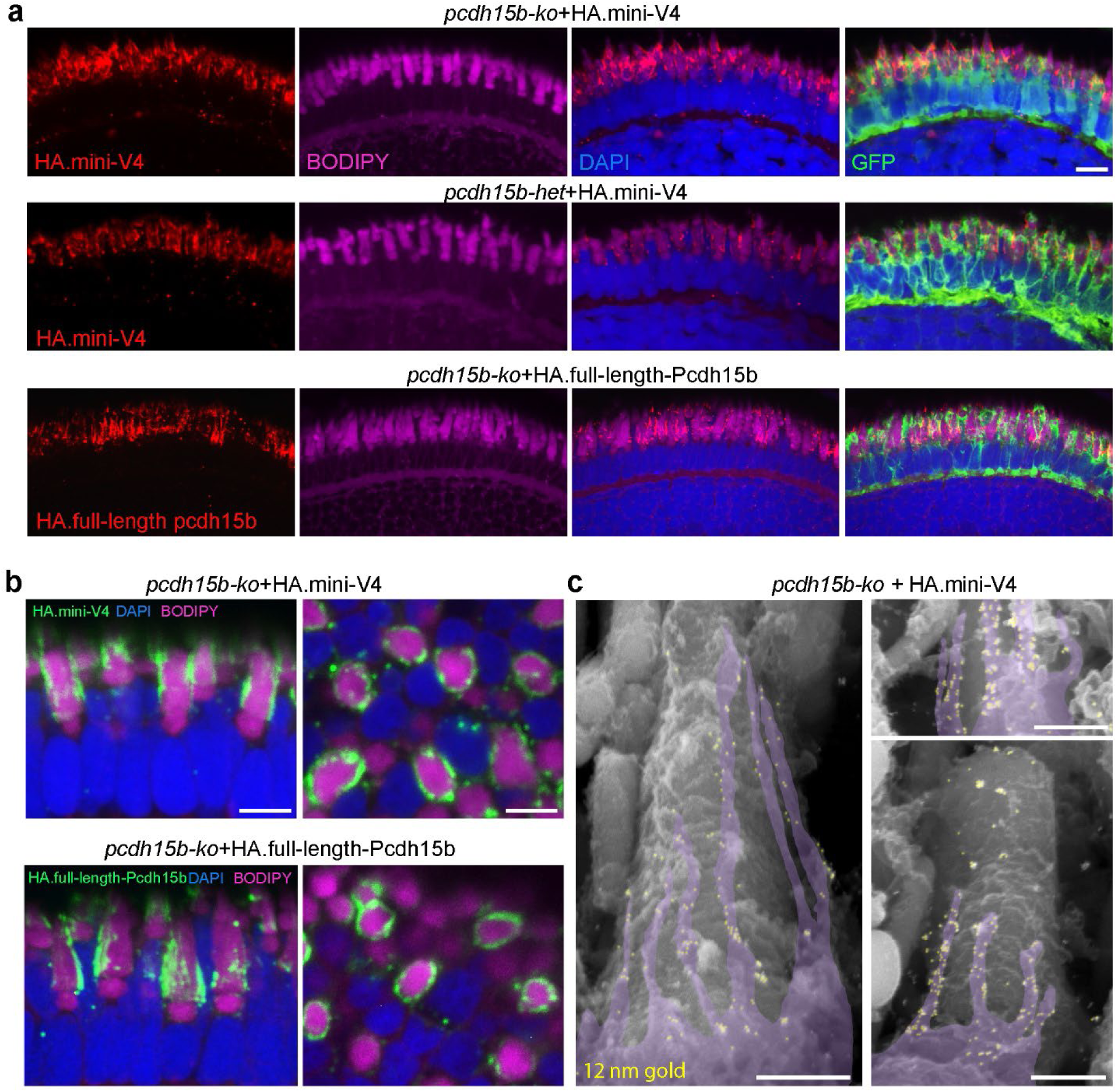
Mini-V4 localizes similarly to full-length Pcdh15b in photoreceptors and supports photoreceptor survival in pcdh15b knockout retina. **a** Confocal images of 7 dpf zebrafish retinas expressing HA.mini-V4 in pcdh15b knockout or heterozygous backgrounds or HA-tagged full-length Pcdh15b. Both constructs showed enrichment at the apical region of photoreceptors, overlapping with BODIPY-labeled outer segments. **b** Higher-magnification views revealed HA-tag immunoreactivity distributed along the OS membranes of photoreceptors (left), forming a ring-like structure in the transversal view of photoreceptors (right). Localization patterns of mini-V4 closely resembled those of the full-length construct, indicating preserved targeting of the mini-Pcdh15b. **c** Immunogold scanning electron micrographs demonstrate proper localization of gold beads labeled mini-V4 (yellow) along calyceal processes (purple). Scale bars: **a** 10 μm, **b** 5 μm, **c** 0.5 μm.

To further verify localization at the ultrastructural level, we performed immunogold scanning electron microscopy on *pcdh15b-ko* retinas expressing mini-V4 (**Figure 7c**). Gold beads marking HA.mini-V4 were concentrated along the calyceal processes that encircle the photoreceptor OS, confirming precise subcellular targeting.

To assess whether mini-V4 maintains proper subcellular targeting in mature photoreceptors, we examined adult *pcdh15b-wt* retinas by confocal immunofluorescence (**Figure 8**). In wild-type retinas, endogenous Pcdh15b localized prominently to the calyceal processes and OS membranes, as shown by strong anti-Pcdh15 labeling overlapping with BODIPY-stained OS (**Figure 8a**). Targeted expression of HA-tagged mini-V4 or full-length Pcdh15b in transgenic *pcdh15b-wt* retinas produced an anti-HA localization pattern indistinguishable from that of the endogenous anti-Pcdh15 pattern (**Figure 8b, c**). In both cases, anti-HA immunoreactivity was concentrated at the apical surfaces of photoreceptors and along their calyceal processes. Higher-magnification imaging confirmed that both constructs extended along microvilli-like projections and OS membranes.

**Figure 8.**
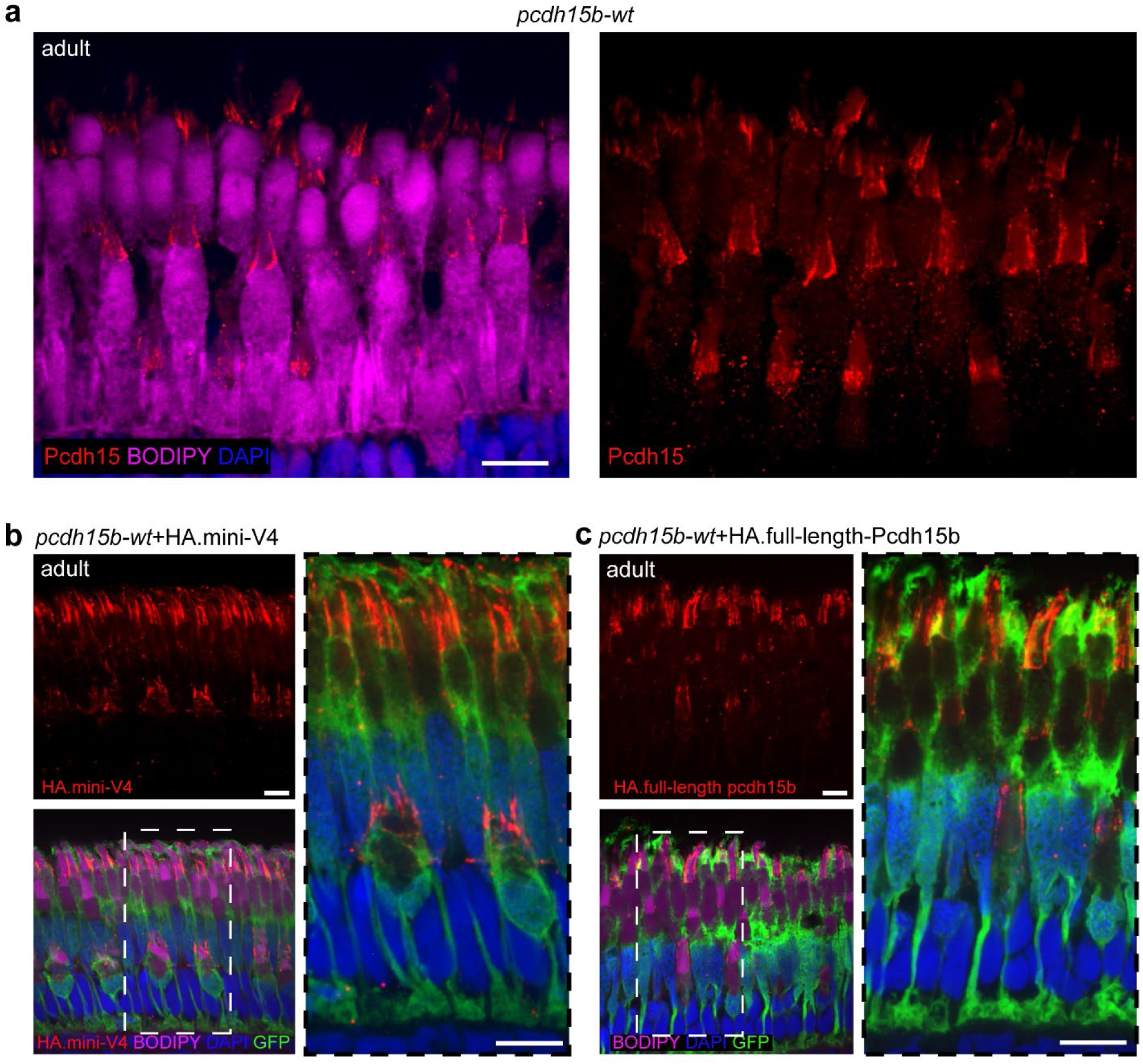
Both mini-V4 and full-length Pcdh15b exhibit subcellular localization patterns in the adult retina consistent with endogenous Pcdh15b localization. **a** Immunostaining of adult *pcdh15b* wild-type retina with anti-Pcdh15 (red) revealed strong labeling along the calyceal processes and OS membranes of photoreceptors. BODIPY (magenta) marks OS membranes, and DAPI (blue) labels nuclei. Left: merged channels; right: Pcdh15 channel only. **b, c** Expression patterns of HA-tagged mini-V4 (**b**) and full-length Pcdh15b transgenes (**c**) in adult *pcdh15b-wt* retina. **b** Top left: anti-HA staining (red) of HA.mini-V4; bottom left: merged view showing HA.mini-V4 (red), BODIPY (magenta), GFP (green) expressed from the transgene, and DAPI (blue). Right: higher magnification of boxed region highlighting HA.mini-V4 localization to calyceal processes and OS. **c** Top left: anti-HA staining (red) of HA.full-length-Pcdh15b; bottom left: merged view showing full-length Pcdh15b (red), BODIPY (magenta), GFP (green), and DAPI (blue). Right: higher magnification of boxed region showing full-length Pcdh15b localization similar to mini-V4. Both constructs targeted correctly to the calyceal processes and OS membranes, mirroring the endogenous Pcdh15b pattern, indicating that mini-V4 retains the proper subcellular localization signals required for function in mature photoreceptors. Scale bars: 10 μm.

These findings demonstrate that mini-V4 faithfully reproduces the subcellular localization of full-length Pcdh15b, ensuring precise targeting to photoreceptor support structures. Moreover, mini-V4 expression effectively preserves photoreceptor architecture and cellularity in *pcdh15b* mutant zebrafish, underscoring its potential as a safe and effective therapeutic approach for retinal degeneration associated with USH1F.

### Mini-V4 expression restores visual behavior and retinal function in *pcdh15b* mutant zebrafish

To evaluate the functional consequences of *pcdh15b* loss and the potential for rescue by mini-V4, we assessed visual performance and retinal function in 7 dpf zebrafish larvae using optokinetic response (OKR) and electroretinography (ERG) assay. In the OKR assay, which measures eye movements elicited by a moving visual pattern (**Figure 9a**), *pcdh15b-ko* larvae displayed significantly reduced numbers of saccades during a 90-second stimulus period compared to wild-type and heterozygous controls (**Figure 9b**), consistent with impaired visual tracking. Moreover, the angle range of individual saccades was significantly restricted in *pcdh15b-ko* larvae (**Figure 9c**), indicating reduced visual field. Expression of mini-V4 in *pcdh15b-ko* larvae robustly improved OKR performance. Saccade number was fully restored to wild-type levels, whereas the angle range of saccades showed partial recovery in mini-V4 expressing *pcdh15b-ko* larvae (**Figure 9b, c**). Importantly, mini-V4 expression in control animals (*pcdh15b*-*wt; -het*) did not adversely affect OKR performance, further supporting the safety of mini-Pcdh15.

**Figure 9:**
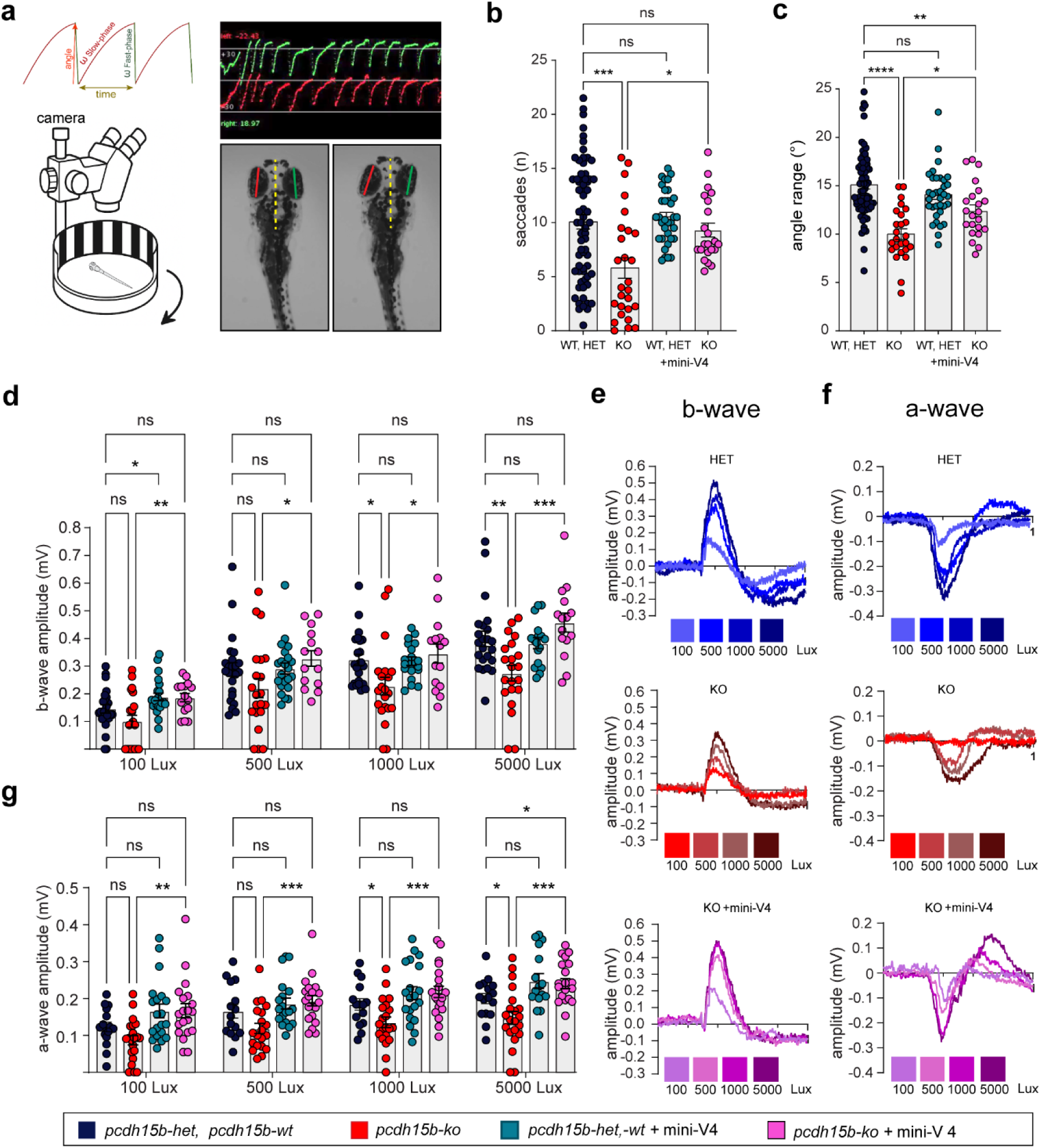
Expression of mini-V4 rescues visual and retinal function in *pcdh15b-ko* zebrafish larvae. **a** Schematic of the optokinetic response (OKR) measurement setup (left). Video camera tracked eye movements in an immobilized larva while a drum with vertical bars was rotated. Example recording traces (right) show sample saccade quantification and angle range. **b** Scatter bar plot of OKR results quantifying the number of saccades during the 90-second recording period demonstrates reduced saccade number in *pcdh15b-ko* (red) compared to *pcdh15b-wt, -het* (blue) and *pcdh15b-wt, -het* with mini-V4 expression (teal), while mini-V4 expression in *pcdh15b-ko* (purple) returned saccade numbers to wild-type level. **c** Scatter bar plot of OKR results quantifying the angle range of individual saccades shows a more limited angle range in *pcdh15b-ko* (red) compared to *pcdh15b-wt; -het* (blue), and mini-V4 expression partly restored the angle range of saccades in *pcdh15b-ko* (purple). **d** Scatter bar plot of electroretinogram (ERG) b-wave amplitudes across light intensities (100, 500, 1000, and 5000 Lux). **e** Representative ERG b-wave traces at increasing intensities for three genotypes (blue, *pcdh15b-wt;-het*; red, *pcdh15b*-*ko*; purple, *pcdh15b*-*ko* with mini-V4 expression). **f** Representative ERG a-wave traces at the same light intensities for genotypes shown in (**e**). **g** Quantification of ERG a-wave amplitudes across four light intensities for genotypes shown in (**b**). Statistical analyses in (**b**-**d**) and (**g**) were performed using ANOVA. ns, not significant; \**P*≤0.05, \*\**P*≤0.01, \*\*\**P*≤0.001, \*\*\*\**P*≤0.0001.

To assess retinal function in *pcdh15b* mutant and rescued larvae directly, we performed electroretinography (ERG) at 7 dpf, recording the millivolt electrical signal at the cornea generated by the retinal response to light. In the vertebrate retina, cones develop before rods, and in zebrafish, cone photoreceptors become functional as early as 4 dpf (Chrispell, Rebrik, & Weiss, 2015; Fadool & Dowling, 2008; Harada, Harada, & Parada, 2007; Schmitt & Dowling, 1999), whereas rod-mediated ERG responses emerge later, between 11 and 21 dpf (Bilotta, Saszik, & Sutherland, 2001). Thus, the 4–7 dpf zebrafish retina function as an all-cone system, making it ideal for studying cone-specific visual responses. However, the ERG signal is dominated by the b-wave, which reflects the activity of ON bipolar cells—postsynaptic to the photoreceptors—and can obscure direct analysis of phototransduction. We first analyzed the slope of the b-wave rising phase across a range of light intensities. In wild-type larvae, the slope increased linearly with the logarithm of light intensity (**Figure 9d, e**). In contrast, *pcdh15b-ko* larvae showed markedly slower b-waves and significantly reduced amplitudes at higher light levels (1000 and 5000 lux) (**Figure 9d, e**), consistent with impaired signal transmission from photoreceptors to ON bipolar cells. These data suggest that phototransduction or synaptic transmission downstream of phototransduction is compromised in *pcdh15b*-deficient retinas. Notably, expression of mini-V4 in *pcdh15b-ko* animals fully restored b-wave amplitudes across all light intensities. No adverse effects or signs of toxicity were observed when mini-V4 was expressed in wild-type or heterozygous backgrounds.

Because the native a-wave is largely masked by the b-wave under standard conditions, we could not directly evaluate photoreceptor function in larvae. To isolate the cone-driven a-wave, which reflects the photoreceptor mass receptor potential, we used a pharmacological approach. An additional group of larvae was pretreated with 500 μM L-(+)-2-amino-4-phosphonobutyric acid (L-AP4), a selective mGluR6 receptor agonist that blocks ON-bipolar cell signaling and effectively eliminates the b-wave. Larvae were immersed in L-AP4 for 5 minutes prior to placement on a PVA sponge for ERG recordings, following previously described protocols (Chrispell, Rebrik, & Weiss, 2015; Lin, Vollrath, Mangosing, Shen, Cardenas, & Corey, 2016; Nelson & Singla, 2009). All subsequent procedures remained unchanged. A-wave amplitudes were significantly reduced in *pcdh15b-ko* larvae, consistent with impaired cone phototransduction, and were fully restored following mini-V4 expression (**Figure 9f, g**).

Together, these findings demonstrate that mini-V4 effectively restores both visual behavior and cone-mediated retinal responses in *pcdh15b* mutant zebrafish, highlighting its therapeutic potential for treating retinal dysfunction.

## DISCUSSION

In this study, we further validated a zebrafish *pcdh15b* mutant model that faithfully reproduces the structural and functional visual defects associated with USH1F (Phillips et al., 2023). The mutants displayed profound disruptions in photoreceptor architecture, including disorganized and shortened OS, abnormal calyceal processes, and impaired visual behavior and retinal function. These phenotypes were exacerbated by bright light exposure, consistent with previous observations in zebrafish and *Xenopus* models of USH1 (Miles et al., 2021; Phillips et al., 2023; Schietroma et al., 2017). Importantly, targeted expression of a zebrafish mini-V4 construct, equivalent to that previously shown to restore hearing in USH1F mouse models (Ivanchenko et al., 2023b), rescued photoreceptor structure, restored calyceal process morphology, and improved both behavioral and electrophysiological measures of visual function. Our findings extend earlier work demonstrating that PCDH15 is essential for the integrity of calyceal processes and OS organization in species that possess these structures. Unlike mice, zebrafish photoreceptors exhibit robust Pcdh15b expression at the calyceal processes along with early, severe deficits to photoreceptor morphology and function, making them a relevant model for studying USH1F retinopathy.

Moreover, the improvement in visual function among mini-V4–treated mutants correlated with enhanced survival beyond the critical second week post-fertilization, when yolk reserves are depleted and active prey tracking becomes essential. Untreated *pcdh15b* mutants exhibit vestibular dysfunction and impaired visual behavior that likely compromise feeding and lead to high lethality after 10 dpf. In contrast, a subset of mini-V4–treated larvae survived into juvenile and adult stages, likely due to restored cone-mediated visual tracking that enhances feeding efficiency despite residual vestibular impairment. These findings highlight the physiological relevance of visual rescue in promoting overall survival and growth in the absence of systemic or vestibular correction.

At the cellular level, mini-V4 not only localized correctly to the calyceal processes and OS membranes but also re-established their structural organization. The restoration of both the number and arrangement of calyceal processes indicates that the mini construct retains essential extracellular cadherin domains for forming and stabilizing these actin-rich microvilli. Correspondingly, the observed improvement in photoreceptor survival and OS integrity suggests that PCDH15-mediated adhesion at calyceal processes is critical for maintaining mechanical stability and supporting proper disc morphogenesis.

Functionally, mini-V4 expression restored visual tracking behavior and improved both b-wave and a-wave ERG responses, indicating rescue of synaptic transmission and/or photoreceptor function. The absence of adverse effects in wild-type or heterozygous animals further supports the safety of the construct in the context of cone-specific expression.

Overall, these results demonstrate that cone-targeted expression of mini-V4 effectively restores both photoreceptor structure and function in *pcdh15b*-deficient zebrafish, highlighting the translational potential of mini-PCDH15 gene therapy for treating USH1F-associated blindness. Future studies should further evaluate the long-term safety and efficacy of this approach and investigate its feasibility for visual rescue in larger, diurnal mammalian models, including non-human primates.

## METHODS

### Zebrafish strains and maintenance

All zebrafish experiment protocols were approved by Harvard University’s Institutional Animal Care and Use Committee, protocol # IS00002804-6. Adult zebrafish were housed at 28.5℃ on a 12-hour light/12-hour dark cycle in 2.8L tanks accommodating up to 12 fish per tank. They were fed twice daily during the light cycle, with at least 4 hours between feedings. Adult fish, age 3 months to 1.5 years, were crossed to produce embryos and larvae. Embryos were then collected, cleaned, and maintained in Embryo Media at about 60 embryos per petri dish in a 28.5℃ incubator. All experiments were performed between 6 dpf and 7 dpf, and all experimental larvae were euthanized by immersion in ice water for 15 minutes, followed by immersion in bleach (1:5 dilutions) either at the end of the experiments or before 8 dpf. Prior to 7 dpf, zebrafish larvae do not undergo gender differentiation, and juveniles and adults used for analyses included equal numbers of males and females.

Zebrafish (*Danio rerio*) of the ABC wild-type strain were used to generate transgenic lines. The *pcdh15b* and *pcdh15a* mutant lines were originally generated at the University of Oregon, Eugene to model the pathogenic R245X mutation that truncates in the EC2 domain. The *pcdh15b* mutant harbors a 7-bp deletion in exon 8, resulting in a frameshift and premature stop codon (Phillips et al., 2023). The *pcdh15a* mutant carries a 20-bp deletion in exon 9, also causing a frameshift and premature stop codon that disrupts all known *pcdh15a* transcript variants. Double mutants (*pcdh15a;pcdh15b*) were generated by crossing heterozygous *pcdh15a* and *pcdh15b* knockout animals.

To generate stable transgenic lines expressing either mini-HA.Pcdh15b-V4 or full-length-HA.Pcdh15b, embryos from heterozygous *pcdh15b* adults were collected and injected at the one-cell stage with Tol2 transposase mRNA (650 ng/μL) along with a bicistronic plasmid (1500 ng/μL). The constructs—Gnat2-HA.mini-pcdh15b-V4-Gnat2-eGFP or Gnat2-HA.full-length-pcdh15b-Gnat2-eGFP—drove expression under zebrafish cone-specific *Gnat2* promoter using a Tol2-based transposon insertion strategy.

Injected mosaic animals were raised to adulthood and crossed with wild-type fish to identify founders with germline transmission, confirmed by fin-clip genotyping. Larval zebrafish from the F2 and subsequent generations were used for all subsequent experiments.

### Adult and Larval Zebrafish Genotyping

All larvae used in this study were obtained from in-crosses of verified heterozygous male and female adult zebrafish, resulting in progeny with three possible genotypes for each mutated gene: wild type, heterozygous, and homozygous. To determine adult genotypes, fish were anesthetized using 0.02% tricaine buffered with sodium bicarbonate, and a small section of the anal fin was collected for DNA extraction and genotyping. To determine larval genotypes, larvae were euthanized by freezing after experiments, and the entire larva was used for DNA extraction. Tissues were digested using a 1:100 dilution of Proteinase K in DirectPCR Tail buffer, and the purified DNA was subjected to PCR amplification of the target gene.

For genotyping the *pcdh15b* allele, the following primers were used: Forward: 5′-TCTGATAGTGGACCAAATATTGC-3′, Reverse: 5′-AGAACTATGTTCAAATGCAAACC-3′ Because the *pcdh15b* knockout zebrafish differ from wild type by only a 7-bp deletion, standard gel electrophoresis alone cannot distinguish genotypes. Therefore, an additional restriction digestion step using BstNI (Bstl-v2), which targets a site present only in the wild-type allele, was performed. After digestion, PCR products were separated on a 1.5% agarose gel. Wild-type alleles produced two bands (390 bp and 343 bp), heterozygotes displayed three bands (733 bp, 390 bp, and 343 bp), and homozygous mutants showed a single uncut band of 733 bp.

For *pcdh15a* genotyping, the following primers were used: Forward: 5′- CCGAAGGTTGTATCTAAAGGTTTGT-3′, Reverse: 5′-GGTACGTGACGGGGTTACAG-3′. As the *pcdh15a* mutant contains a 20-bp deletion that is not easily resolved by standard electrophoresis, a restriction digestion with AvaII was required. The wild-type allele contains an AvaII site and yields two bands (∼142 bp and 66 bp), while the mutant allele lacks the site and yields a single 188 bp band. Heterozygotes show all three bands. PCR products were resolved on a 1.5% agarose gel.

To confirm the presence of injected transgenes (mini-*pcdh15b*-V4 or full-length *pcdh15b*), an additional PCR was performed using the following primers: Forward: 5′- ACCTGGTGATCAAGGCAGAG-3′, Reverse: 5′-CAGCATGTACAGCGGTCACT-3′. Samples lacking the transgene showed no bands. The expected product size was approximately 174 bp for mini-*PCDH15*-V4 and 1770 bp for the full-length *PCDH15*. Products were visualized on a 1.5% agarose gel.

### Light Exposure Procedure

To exacerbate and assess the retinopathy phenotype in larval zebrafish, a 72-hour light exposure procedure was implemented. Larvae kept in transparent Petri dishes were moved into a custom incubator with an overhead LED light at 3 dpf for LED light exposure protocol (12-hour light/12-hour dark, 3000 lux). At the end of 6 dpf, larvae were moved to the dark for overnight acclimation before electroretinogram (ERG) or optokinetic response (OKR) testing.

### Electroretinogram (ERG)

On the day of ERG experiments, larvae were tested individually, and all procedures were completed under red illumination only to minimize bleaching of the visual pigment. For regular ERG experiments testing both the initial a-wave from photoreceptor activity and the larger b-wave originating from ON bipolar cell depolarization, single larvae were removed from the Petri dish and positioned laterally on an embryo media-soaked PVA sponge with an embedded reference electrode. Larvae were always placed on their right side for ERG testing of the left eye. The recording electrode wire was inserted into the embryo media-filled recording pipette, which was then placed carefully on the cornea with slight pressure. Light flashes of five different light intensities (50 lux, 100 lux, 500 lux, 1000 lux, 5000 lux) were tested for each larva, with each intensity measured consecutively three times with 200 ms intervals and a 1-minute wait time between each light intensity level. An additional group of larvae was tested for a-wave only (Chrispell, Rebrik, & Weiss, 2015; Lin et al., 2016; Nelson & Singla, 2009). To block the b-wave, larvae were immersed in a 500 μM solution of 2-amino-4-phosphonobutyric acid, a type III metabotropic agonist, for 5 minutes before they were positioned on the PVA sponge. All subsequent steps remained the same. Data were recorded using Clampex and analyzed with Clampfit (Molecular Devices). At each light stimulation intensity, results from the three consecutive recordings were averaged.

### Optokinetic response (OKR)

On the day of OKR experiments, 7 dpf larvae were tested individually, and all procedures were completed under red illumination only. A single larva was transferred from its home Petri dish to a drop of 2% methylcellulose on a 35 mm Petri dish. The larva was carefully positioned with the dorsal side up and the eyes equidistant from the surface. The methylcellulose was spread outward to limit larval movements. The larva was allowed 20 minutes of methylcellulose acclimation and was then moved into the OKR drum for testing. The cylindrical OKR drum was 10 cm in diameter and 3 cm in height, with vertical alternating 1.5 cm wide black and white stripes. Two 90-second episodes of OKR were recorded for each larva: clockwise and counterclockwise OKR drum rotations, with 180 seconds of acclimation in between. The overhead camera captured larval eye movement at 25 frames per second, and all frames were further analyzed in FIJI and MATLAB to measure angular eye deflection.

### Immunofluorescence labeling of zebrafish

Larval zebrafish were euthanized with an ice bath. Tails were cut and digested for genotyping, while heads were collected and fixed overnight at 4℃ in 4% formaldehyde in Hanks’ Balanced Salt Solution (HBSS) with calcium and magnesium. After three 10-minute rinses with HBSS, the samples were cryoprotected with an ascending concentration series of sucrose solutions (10%, 20% and 30%). The samples were then embedded in the OCT compound and stored at −80°C. Cryosections were generated using a Leica CM 3050 S cryostat at 20-μm step size.

For immunofluorescence labeling, the following primary and secondary antibodies were used: anti-PCDH15 antibody (R&D Systems, AF6729), rabbit anti-HA (C29F4) antibody (Cell Signaling Technology, 3724), mouse monoclonal anti-rhodopsin (1:500) (MilliporeSigma, MAB5316), donkey anti-rabbit IgG secondary antibody conjugated to Alexa Fluor 594 (1:200) (Invitrogen, R37119), donkey anti-sheep IgG conjugated to Alexa Fluor 488 (1:200) (Invitrogen, A-11015), donkey anti-mouse IgG conjugated to Alexa Fluor 488 (1:200) (Invitrogen, A32766), donkey anti-mouse IgG conjugated to Alexa Fluor 405 (1:200) (Invitrogen, A48257), and donkey anti-rabbit IgG conjugated to Alexa Fluor 488 (1:200) (Invitrogen, A-21206).

Before staining, slides were brought to room temperature and were blocked with 10% donkey serum in PBS for 1 hr. Primary antibodies were diluted in 10% donkey serum and incubated overnight at room temperature, followed by three 10-minute rinses in PBS. Next, samples were incubated in 10% donkey serum for 1 hr and then incubated overnight at room temperature with a secondary antibody in a 1:200 dilution in 10% donkey serum, the blocking solution. We used DAPI to label cell nuclei (1:500), BODIPY to label membranes (1:1500), and phalloidin to label actin-rich calyceal processes. Tissues were mounted on a Colorfrost glass slide (Thermo Fisher Scientific) using Prolong Gold Antifade mounting medium (Thermo Fisher Scientific). Imaging was performed with a Nikon Ti2 inverted spinning disk confocal using a Plan Fluor 40×/1.3 oil objective, Plan Apo λ 60×/1.4 oil objective, and a Plan Apo λ 100×/1.45 oil objective.

### Transmission electron microscopy

Larvae of 7 dpf were euthanized with an ice bath and fixed with 2.5% glutaraldehyde in 0.1 M cacodylate buffer for 24 hours at 4℃. After three rinses in cacodylate buffer, samples underwent 24-hour postfixation with 1% osmium tetroxide/1.5% potassium ferrocyanide in 0.1 M cacodylate buffer at room temperature in the dark. Then, samples were washed three times in 0.1 M cacodylate buffer (pH 7.2), briefly rinsed in distilled water, dehydrated in an ascending series of ethanol concentrations, equilibrated in propylene oxide, and finally infiltrated and embedded in epoxy resin (Araldite 502/Embed-812 embedding media). The resin was allowed to polymerize at 60°C for 48 hours, and the resin blocks were then sectioned at 60–80 nm intervals using a Reichert Ultracut S ultramicrotome. Sections were mounted on copper Formvar/Carbon-coated grids and examined with a JEOL 1200EX microscope operating at 80 kV.

### Scanning electron microscopy

The retinal pigment epithelium and neural retina from enucleated eyes were quickly dissected apart and fixed with 2.5% glutaraldehyde in 0.1 M cacodylate buffer (pH 7.2) for 1-2 hours at room temperature. After three rinses in cacodylate buffer, samples underwent 4-hour postfixation with 1% osmium tetroxide in 0.1 M cacodylate buffer at room temperature in the dark. Samples were then rinsed with 0.1 M cacodylate buffer (pH 7.2) and distilled water, dehydrated using an ascending ethanol series, and critically dried with liquid CO_2_ (Tousimis Autosamdri 815). Subsequently, the samples were mounted on aluminum stubs with carbon conductive tabs, sputter-coated (Leica EM ACE600 sputter coater) with 5-nm platinum, and examined using a field-emission scanning electron microscope (Hitachi S-4700).

### Immunogold scanning electron microscopy

Immunogold scanning electron microscopy was performed following previously described protocols (Ivanchenko et al., 2024; Ivanchenko, Indzhykulian, & Corey, 2021). Larval retinas were fixed in 4% formaldehyde, then washed and blocked for 2 hours at room temperature with 10% goat serum. Next, samples were incubated overnight at room temperature with a primary anti–HA tag antibody (rabbit anti-HA (C29F4) antibody (Cell Signaling Technology, 3724; 1:200 in 10% goat serum) and rinsed in HBSS. Following an additional 30-minute block with 10% goat serum, samples were incubated overnight at room temperature with a secondary antibody—12 nm Colloidal Gold AffiniPure Goat Anti-Rabbit IgG (Jackson ImmunoResearch, 111-205-144; 1:30 in blocking solution). After secondary labeling, samples were rinsed in HBSS. Next, they were fixed with 2.5% glutaraldehyde in 0.1 M cacodylate buffer (pH 7.2) for 1-2 hours at room temperature. After three rinses in cacodylate buffer and one in distilled water, they were dehydrated using an ascending ethanol series, and critically dried with liquid CO_2_ (Tousimis Autosamdri 815). Subsequently, the samples were mounted on aluminum stubs with carbon conductive tabs, sputter-coated (Leica EM ACE600 sputter coater) with 5-nm palladium, and examined using a back-scattered detector in a Hitachi S-4700 scanning electron microscope.

### RNA extraction, cDNA production, reverse transcription, PCR amplification, and sequencing

Larval zebrafish at 7 days post-fertilization (dpf) were euthanized by rapid cooling on ice. Eye globes were dissected and collected, while the remaining larval tissue was used for DNA extraction and genotyping. Eyes were stored in Eppendorf tubes until genotyping was completed. Subsequently, tubes were thawed on ice, and 10 eyes from each genotype were pooled together. Each pooled sample was incubated with 50 μL of 2× digestion buffer and 10 μL of Proteinase K (6 U) for 10 min at room temperature, with occasional gentle pipetting to facilitate tissue dissociation. Following digestion, 60 μL of RNA lysis buffer (equal to the digestion mixture volume) was added, and samples were incubated for 5 min at room temperature. Lysates were then centrifuged at 12,000 × g for 5 min, and the supernatants were collected; pellets were discarded.

Total RNA was purified using the Quick-RNA Microprep Kit (Zymo Research, #R1050) according to the manufacturer’s instructions. Reverse transcription was performed using the SuperScript IV VILO Master Mix with ezDNase enzyme (Invitrogen, #11766050) to generate cDNA. Pcdh15b cDNA was amplified using a forward primer in 5′-GGACATTCGACATCCCCCTC-3′ and a reverse primer 5′-CTGCAAGGCAGAAACATCGG-3′, yielding an expected 202-bp product. Pcdh15a cDNA was amplified using a forward primer 5′- ACAATGGAGCCACGGACATT-3′ and a reverse primer in 5′-GATGGGAGGCGTCACATTCA-3′, yielding an expected 395 bp product.

β-actin was amplified using the primers 5′-AAGCAGGAGTACGATGAGTC-3′and 5′-TGGAGTCCTCAGATGCATTTG-3′, producing a 238-bp fragment. PCR products were gel-purified using the DNA Gel Extraction Kit (Monarch, #T1020S). Purified products were subcloned, transformed into competent cells, mini-prepped, and Sanger sequenced using the NEB PCR Cloning Kit (NEB, #E1202S).

## ACKNOWLEDGEMENTS

We appreciate the use of the Nikon Ti2 inverted spinning disk confocal and Olympus VS200 Slide Scanner at the Harvard Medical School MicRoN Microscopy Core, use of the Hitachi S-4700 scanning electron microscope at the Harvard Medical School Electron Microscopy Facility, and use of instruments at the Harvard University Center for Nanoscale Systems. We would like to thank Dr. Artur A. Indzhykulian for his valuable guidance on ERG methodology and construct design. This work was supported by a Bertarelli Foundation, Usher 1F Collaborative, Blavatnik Therapeutic Challenge Award, and Foundation Fighting Blindness.

## AUTHOR CONTRIBUTIONS STATEMENT

M.V.I., X.C., D.M.H., A.K., K.T.A.B, M.H., A.L., B.H.D., C.G., data acquisition, data analysis, data interpretation, visualization. J.B.P., J.W., and M.W. generated and provided zebrafish models and contributed to data interpretation. A.R.M. and S.G.M. offered guidance on methodology, including construct design and data analysis. D.P.C. and M.V.I. conceived and supervised the study and wrote the manuscript with input from all authors.

---

**Supplementary Figure 1:**
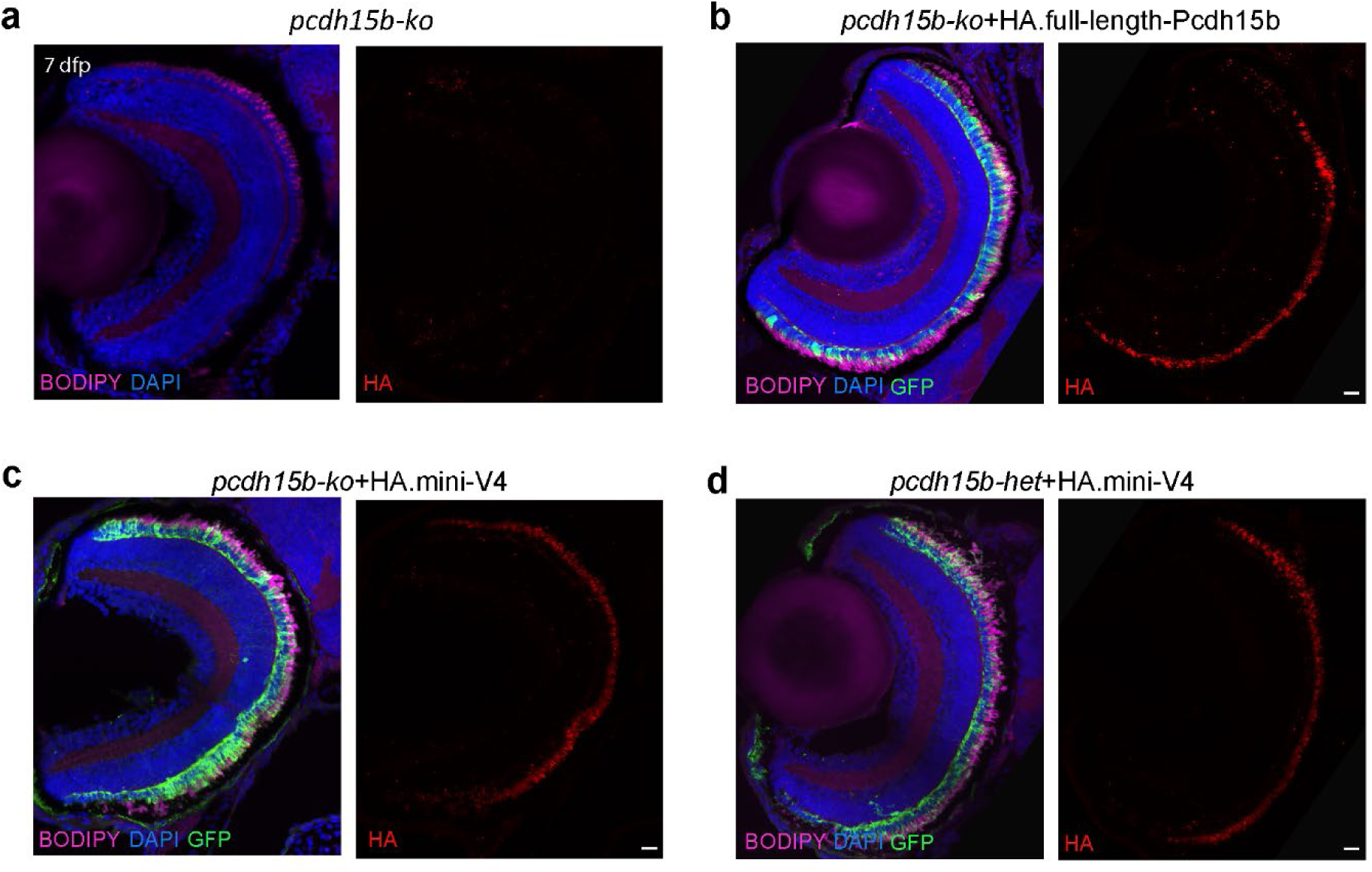
Mini-V4 localizes to photoreceptors in the zebrafish retina similarly to full-length Pcdh15b. a-d. Confocal images of 7 dpf zebrafish eyes showing transverse retinal sections stained with anti-HA (red), BODIPY (magenta), DAPI (blue), and GFP (green). **a** A negative control, *pcdh15b-ko* larvae without any transgene, showed no HA signal. **b** HA-tagged full-length Pcdh15b in the *pcdh15b-ko* background showed a strong HA signal localized to the photoreceptor layer. **c** HA-tagged mini-V4 in the *pcdh15b-ko* background exhibited a similar localization pattern to full-length Pcdh15b, with enrichment in the photoreceptor layer. **d** In the *pcdh15b-het* background, mini-V4 also showed a robust HA signal in photoreceptors with no signs of toxicity. These data confirm that mini-V4 recapitulates the subcellular distribution of full-length Pcdh15b in the retina. Scale bars: 10 μm.

